# When do shifts in trait dynamics precede population declines?

**DOI:** 10.1101/424671

**Authors:** Gaurav Baruah, Christopher F. Clements, Fréderic Guillaume, Arpat Ozgul

**Affiliations:** Department of Evolutionary Biology and Environmental studies, University of Zurich; School of Biosciences, University of Melbourne

**Keywords:** population decline, quantitative trait, early warning signals, demography

## Abstract

Predicting population responses to environmental change is an on-going challenge in ecology. Studies investigating the links between fitness-related phenotypic traits and demography have shown that trait dynamic responses to environmental change can sometimes precede population dynamic responses, and thus, can be used as an early warning signal. However, it is still unknown under which ecological and evolutionary circumstances, shifts in fitness-related traits can precede population responses to environmental perturbation. Here, we take a trait-based demographic approach and investigate both trait and population dynamics in a density-regulated population in response to a gradual change in the environment. We explore the ecological and evolutionary constraints under which shifts in a fitness-related trait precedes a decline in population size. We show both analytically and with experimental data that under medium-to-slow rate of environmental change, shifts in trait value can precede population decline. We further show the positive influence of environmental predictability, average reproductive rate, plasticity, and genetic variation on shifts in trait dynamics preceding potential population declines. These results still hold under non-constant genetic variation and environmental stochasticity. Our study highlights ecological and evolutionary circumstances under which a fitness-related trait can be used as an early warning signal of an impending population decline.

## 1. Introduction

Exogenous pressure can force complex systems with alternative stable states towards so-called tipping points, the point at which the state system can rapidly and substantially change in response to a small perturbation (May 1977; Hutchings and Reynolds 2004; Frank et al. 2011). Examples of such transitions are documented from rapid shifts in shrub cover in grasslands (Kéfi et al. 2007) to the collapse of fisheries (Jackson et al. 2001; Frank et al. 2011), and have been shown to be experimentally inducible in laboratory systems (Dai et al. 2012; Lei Dai, Korolev, and Gore 2013). The non-linear nature of such transitions makes them difficult to predict, but may be possible through the identification of statistical signals embedded in time series data, typically termed early warning signals (EWS) (Wissel 1984; Dakos et al. 2008; Marten Scheffer et al. 2009). Such statistical signals arise from a systems behavior prior to a critical transition (van Nes and Scheffer 2007; M. Scheffer et al. 2012) – whereby it takes longer to return to it’s original equilibrium state after every perturbation. This behavior of the system is known as critical slowing down (CSD).

CSD behavior is predicted to lead to increasing variance and autocorrelation in the abundance time-series in the region of a bifurcation. However, shifts in variance or autocorrelation in the abundance time series might not be the only indicators of whether a population is nearing a tipping point (Clements and Ozgul 2016). External environmental change has been shown to also substantially affect phenotypic trait distributions along with changes in demography (Traill, Schindler, and Coulson 2014; Pigeon et al. 2017). Phenotypic traits, for example body size, are intimately linked with an individual survival and reproductive success (McNamara & Houston, 2007), vulnerability of a population to extinction and fluctuations in population size (Olden, Hogan, and Zanden 2007; van Benthem et al. 2017). Changes in body size have also been shown to influence resilience of food webs to disturbance (Woodward et al. 2005). Furthermore, recent work has also suggested the inclusion of body size-based measures of stability, as shifts in body size of diatoms were detected before a regime shift in a lake ecosystem (Spanbauer et al. 2016). Thus, tracking shifts in such fitness-related phenotypic trait values might allow us to infer how stable the population is, which can inform us about an impending population decline.

Empirical observations from experiments and natural populations have shown that including information from shifts in phenotypic traits such as body size into abundance-based stability indicators improves the accuracy of predictions of population collapse (Clements and Ozgul 2016; Clements et al. 2017), but the circumstances under which doing so is informative is still unknown. The response of a population to a change in the environment can be a combination of ecological and evolutionary responses governed by factors such as genetic variation or adaptive plasticity (Charmantier et al. 2008; Ozgul et al. 2012). Plastic responses to a change in the environment can be fast, and such responses have been shown to stabilize population dynamics (Reed et al. 2010; Charmantier et al. 2008). However, if the environment keeps on changing, populations might deplete its plastic capacity and standing genetic variation, causing it to eventually decline. This decline in population size will also be dependent on the reproductive rate i.e., whether a population is growing faster or slower (Hutchings et al. 2012; Juan-Jordá et al. 2015). In addition to this, due to directional change in the environment, selection on the trait will act on the standing genetic variation of that trait, and higher the genetic variation faster is the evolutionary response (assuming the phenotype heritable). Such theoretical expectations raise important practical questions: can a shift in phenotypic trait dynamics occur before an eventual decline in population size? Under what circumstances is this possible? If such a shift in trait dynamics occurs, what are the factors that govern the earlier occurrence of this shift?

To answer these questions, we used a combination of theoretical and experimental approaches to understand the circumstances under which information from trait values can be useful to predict a potential population decline. First, we integrated quantitative genetics and population dynamics in a theoretical approach, and showed both analytically and numerically whether, and under what circumstances, shifts in trait dynamics can precede population declines. We then experimentally test our predictions using microcosm data where replicate protist populations were forced to collapse under different environmental perturbations (Clements and Ozgul 2016). Finally, we evaluate, through numerical simulations, how genetic variation, adaptive plasticity, and reproductive rate affect when shifts in trait dynamics precede decline in population size.

## 2. Methods and modeling framework

### 2.1 The model

In our model, we consider a closed population that has non-overlapping generations, and is subjected to density-dependent population regulation. We assume in our population that all individuals experience the same environment; i.e. there is no spatial heterogeneity. Fitness of individuals in the population is determined by a quantitative trait *z* that is under stabilizing selection. Under these assumptions the dynamics of the population and the mean value of the trait can be written as (Chevin and Lande 2010; Gomulkiewicz and Holt 1995)

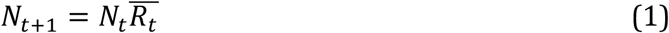

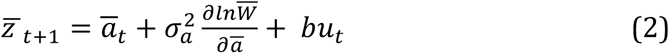

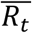 is the average fitness of the population at generation *t*. 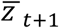 is the mean value of the trait at generation *t+1,ā_t_* is the mean breeding value of the population at generation *t*, 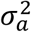 is the additive genetic variance, 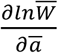 is the gradient of selection on the mean trait 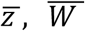 is the mean fitness due to the trait *z* and *bu_t_* quantifies the average plastic response of the trait (Table 1).

**Table 1:** List of variables and parameter values used in the model. Note that variables are not given any value, but parameters for which the model simulations are tested are given values.

| Parameters /variables | Description | Value |
| --- | --- | --- |
| $N_t$ | Abundance of the population at time t | .... |
| $\bar{z}_t ; \bar{a}_t$ | Mean trait value at time t; mean breeding value at time t | .... |
| $\sigma_z^2 ; \sigma_a^2$ | Variance of the mean trait; additive genetic variance or variance of the mean breeding value | [0.05, 0.08, 0.1, 0.2, 0.3, 0.4,0.5, 0.6 ] ;<br>[0.05, 0.08, 0.1, 0.2, 0.3, 0.4,0.5, 0.6 ] |
| $b$ | Strength in phenotypic plasticity | 0.05, 0.1, 0.2,0.3,0.4, 0.5, 0.6 |
| $R_0$ | Net reproductive rate | 1.1, 1.15, 1.2, 1.25, 1.3,1.35, 1.5 |
| $u_t$ | Environmental cue at time t | Case 1: -0.0001<br>Case 2: $-\frac{B}{2} + \frac{B}{2} e^{-\left(\frac{t}{500}C+C\right)}$<br>where, C=4 |
| $\theta_t$ | Optimum trait at time t | .... |
| $w_z^2$ | Width of the Gaussian stabilizing fitness function. A measure of the strength in selection | 40 |
| $z_i, a_i$ | Trait value for an individual i ; breeding value for an individual i | .... |
| $\epsilon$ | Residual component or unexplained component of a trait z | 0 |
| $\sigma_\epsilon^2$ | Variance in that residual component $\epsilon$ | 0 |
| $E_t$ | External environmental value at time t | .... |
| B | Coefficient that captures the strength in the external environmental change | 2 |
| C | Parameter that controls the rate of change in the external environment $E_t$ | Various: 10.5 to 120.5 |

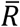 in equation (1) can be expanded as :

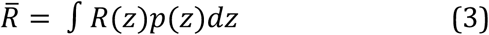

where *R*(*z*) is the absolute multiplicative fitness of an individual with trait *z* and *p*(*z*) is the distribution of the quantitative trait *z* which is normal. *R*(*z*) can further be decomposed into:

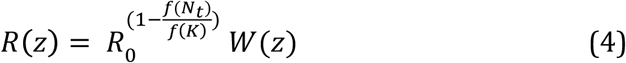

where, *R*_0_ is the net reproductive rate, 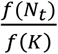 is the density dependence function and *W*(*z*) is the Gaussian stabilizing fitness function given as 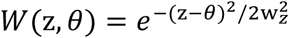 with width 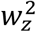 and optimum phenotype of *θ*. Hence an individual’s reproductive fitness will thus be determined not only by how far its trait *z* is from the optimum phenotype *θ*, but will also concurrently be dependent on the current density of the population given by the density-dependence function.

Finally, the response of the primary trait *z* to the environment is modeled using linear reaction norms (Gavrilets and Scheiner 1993). The phenotype of an individual at any generation *t* in the population is given by

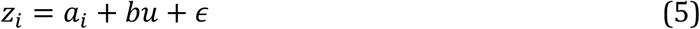

where *a* is the breeding value of the individual *i*, which is normally distributed with mean *ā* and additive genetic variance of 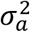. *ϵ* is the residual component with mean 0 and variance 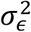. The slope *b* in our model determines how plastic the trait is and is modeled as a constant value meaning that plasticity in the trait cannot evolve. Hence, variance of the phenotype *z* is 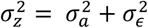 (we assume 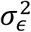 to be zero in our model). The environment in our model determines the optimal phenotypic value *θ* for the primary phenotype *z* and also cues the plastic response. *θ* is assumed to be linearly dependent on the environment *E* that selects for a particular phenotypic value such that at any time *t, θ_t_* = *BE_t_*. The environmental cue *u* quantifies how an individual on average perceives the environment. For example, snow cover could be one of the environmental cues for ground squirrels to come out of their hibernation that correlates with resource availability, the environmental factor (Lane et al. 2012). This means that the cue *u_t_* and the selecting environment *E_t_* are related (Charmantier et al. 2008; Reed et al. 2010). We model this relation by making the environmental cue a function of the selecting environment such that *u_t_* = *f* (*E_t_*). In the case when *u_t_* is a linear function of the selecting environment *E_t_*, changes in the selecting environment triggers a change in the cue response. If *u_t_* is modeled as a constant value and independent of the selecting environment *E_t_*, then the cue response will be different to the change in the selecting environment. In this case, individuals in the population will not be able to perceive changes in the selecting environment.

Finally we can arrive at equation (2), which describes the dynamical changes in the mean trait due to changes in the mean fitness 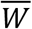 caused by changes in the selecting environment *E_t_*. The mean fitness 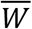 can be calculated by: 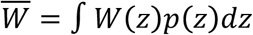 = 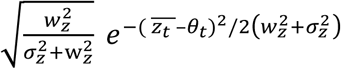

In our model we specifically address questions linked with the decline phase of the population and not evolutionary rescue. Evolutionary rescue is a long-term process, which occurs when genetic evolution rescues a population from extinction in response to changes in the environment. Whether this rescue happens will depend on the initial decline phase of the population, as the population might collapse before it can adapt to changing environmental conditions (Gonzalez et al. 2013). Hence, predicting this initial decline phase is of foremost importance if one has to mitigate the demographic response of the population, before evolutionary rescue takes place at a later phase.

### 2.1 Analytical framework

Optimal phenotypic change in our analyses is directional and given by:

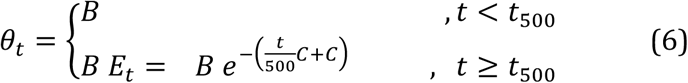

Where *B* = 2, 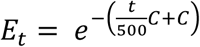, *t* is the time in generations and C is a parameter that controls how fast the optimal phenotype changes over time. We vary C over a range of values to create a gradient of environmental change from fast to slow. Without loss of generality, we consider that the average trait value is at its optimum, i.e. 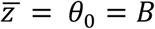 and the environment shifts at t = 500.

Following an environmental change scenario, shifts in trait and population dynamics are calculated by 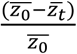 and 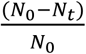 respectively, where 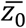 is the initial average trait value and 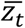 is the average trait value after the optimum phenotype shifts. *N*_0_ is the abundance at carrying capacity and *N_t_* is the abundance after the environment shifts. We consider in the analytical cases below without the effect of plasticity, i.e. *b* = 0 (appendix A2 for plasticity affect). Hence, all the individuals in the population differ only in terms of their breeding value. For analytical simplicity, we show two very simplistic cases of one-generation change in trait and the population after the optimum environment shifts. *Case 1*: the environmental change scenario that causes 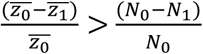 i.e., shift in trait value before population decline and, *Case 2*: the environmental change scenario that causes 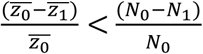 i.e., shift in trait value after a population decline. We consider in the above cases two extreme ends of environmental change: abrupt, large shift in one generation and slow, small shift in one generation. In the case of abrupt and large shift in the environment, the optimum phenotype is allowed to shift in *one generation* by a large magnitude to a new value of *θ*_1_ = 0.5 from a value of *θ*_0_ = 2 such that, 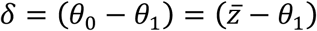 = 1.5. Specifically, *δ* can be termed as the initial phenotypic lag of the mean trait to optimum phenotype in 1 generation. The level of this lag is dependent on how fast the optimum phenotype shifts. A shift of *δ* ≥ 1.5, or a lag of 1.5 in just *one generation* causes a substantial population decline (Fig. 1). Moreover, such a jump in the optimum phenotype to a new value introduces a novel optimum that is beyond the distribution of the adapted trait distribution (with 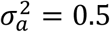). In the case of slow and small shift in the environment, the optimum phenotype in *one generation* is allowed to shift by a very small amount, *δ* ≤ 0.2. This value corresponds to a shift in the optimum that is not novel and within the realms of the adapted trait distribution.

**Figure 1.**
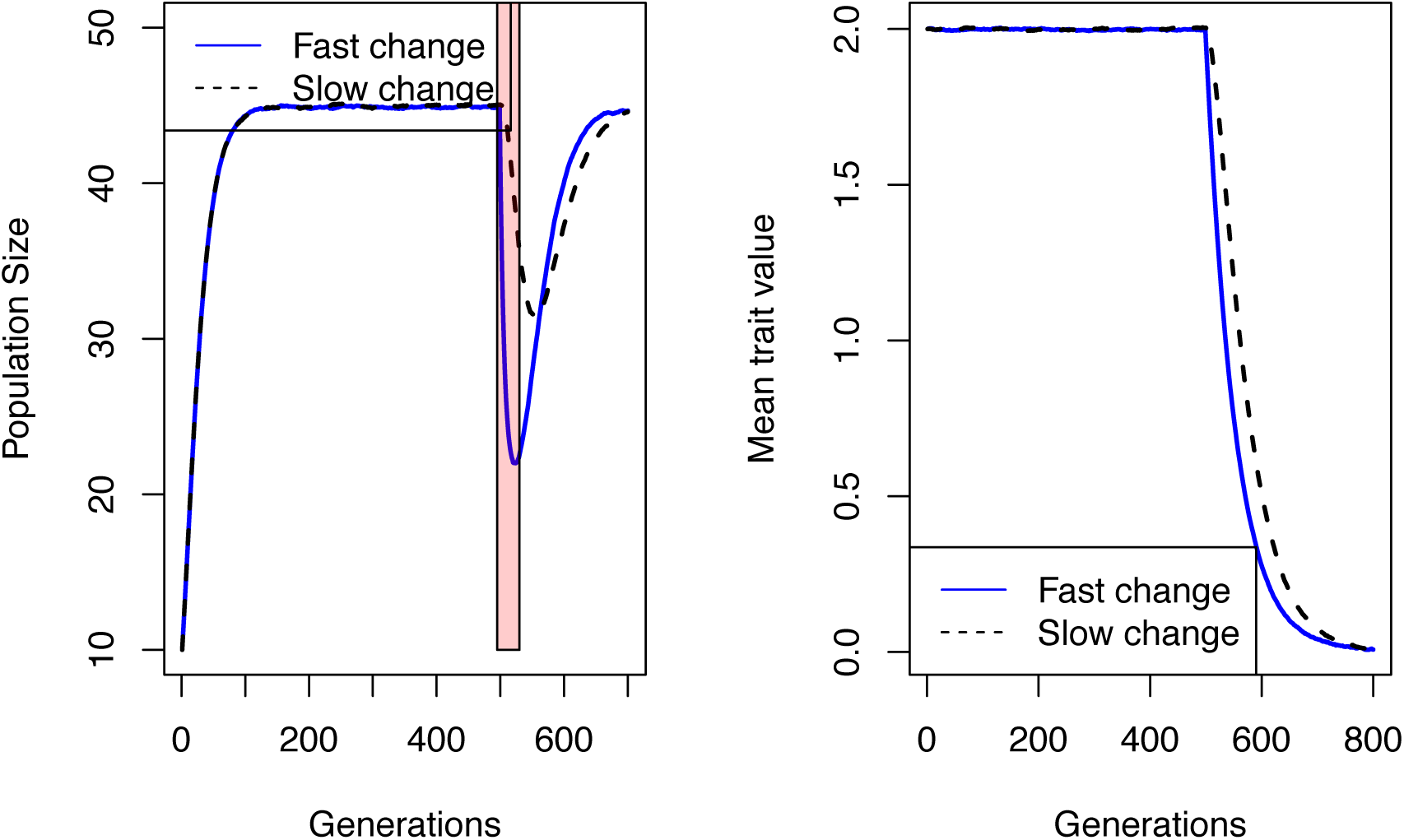
Population and trait dynamics under two different environmental scenarios. Left: Population size declines by ~50%, when the optimum environment shifts fast (C=120.5; blue line), whereas during slow change (C= 10.5; dashed line) population declines but to a lesser extent. Right: Trait dynamics during fast (blue line; C = 120.5) and slow (dashed line; C=10.5) environmental change. Data based on deterministic simulations of the theoretical model. Parameters used: 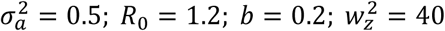.

We discuss the analytical results of case 1 and case 2 in the *Result* section.

### 2.2 Numerical simulations

We performed deterministic numerical simulations of the model described above. We also did stochastic numerical simulations of the model (for details see appendix A3). Dynamics of the trait, population and optimum environment were iteratively updated using equations (1-6). We used Gompertz density function as the form of density-dependence in our model simulations (Chevin and Lande 2010; Gompertz 1825). Without loss of generality, we assumed that the mean trait value is at its optimum 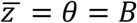 at the start of each simulation. Environmental change was introduced only after the population has reached its carrying capacity which was at t= 500. By varying the parameter C (equation 6), we simulated a range of environmental change scenarios: from very slow (C=10.5) to fast (C= 120.5). In all of our simulations of different rates of environmental change, the optimum environment switched to a new value at *t* = 500. The magnitude of this switch, however, depended on C (equation 6). For example, if C=20.5, the optimum took around ~ 30 generations to switch by a magnitude of 1.5 units and if C=120.5, the optimum took just ~5 generations to switch. Next, to quantify whether the trait or population abundance shifted first when the environmental conditions changed, we calculated the area under the curves (AUC) of both the trait shift 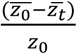 and population shift 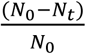 over 20 generations after the optimum environment changed. 1-generation change was used only in the two extreme cases to elucidate the mathematical simplicity behind the environmental change scenario under which trait could shift before a potential population decline. AUC, on the other hand, was used to graphically extend the analytical results of 1-generation change to 20-generation change. AUC quantified both the magnitude and the timing of standardized shifts. With this, we then calculated the metric, *Δshift* = (AUC of shift in mean value of the trait – AUC of population abundance shift). We then compared the size of *Δshift* to the rate of change in the environment (for stochastic change in the environment see appendix A3). If *Δshift* was negative, for a particular rate of change in the environment, it indicated that the population shift was larger and hence the population declined/shifted earlier than a shift in the mean trait value; if *Δshift* was positive it meant a shift in the trait value occurred before a decline/shift in population size. This allowed us to quantify under what environmental change scenarios (from slow to fast) we could expect shift in average trait value to occur before a potential population decline. We then evaluated how the metric *Δshift*, for the range of environmental change scenarios, was affected by factors like genetic variation, strength in plasticity, reproductive rate, and environmental predictability.

#### 2.2.1 Rate of environmental change

We varied the parameter C from a value of 10.5 to 120.5 with a step size of 2 to simulate a gradient of perturbations from slow to fast environmental change. We then calculated from equation (6) the time in generations the optimum took to shift by a magnitude of ~1.5 units. This ~1.5 unit of change in the optimum in 5 generations (when, C=120.5) or in more than ~30 generations (values of *C* ≤ 30) was enough to cause a significant population decline (Fig. 1). We then assessed how *Δshift* was affected by the rate of environmental change.

#### 2.2.2 Environmental predictability

We simulated two specific scenarios: 1) when the cue was a linear function of the optimum environment, and hence individuals of the population could perfectly predict changes in the optimum environment, 2) when the cue was a constant value and hence individuals had zero predictability of changes in the optimum environment. We then assessed how these two scenarios of environmental predictability affected *Δshift*

#### 2.2.3 Genetic variation, strength of adaptive plasticity and average reproductive rate

We ran a range of numerical simulations with different levels of genetic variation (low to high): 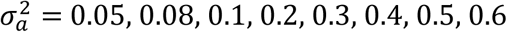; low to high strength of adaptive plasticity b = 0.05, 0.1, 0.2,0.3,0.4, 0.5, 0.6; and low to high average reproductive rate R_0_ = 1.1, 1.15, 1.2, 1.25, 1.3, 1.35, 1.5. While varying levels of genetic variation from low to high, we kept adaptive plasticity strength at b = 0.2 and R_0_ at 1.2. While varying levels of *b*, we kept 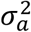 at 0.5 and R_0_ at 1.2. Finally, while varying R_0_, we kept 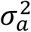 and *b* at 0.5 and 0.2 respectively. For a range of environmental change scenarios, we then evaluated the effect of each parameter value on *Δshift*

#### 2.2.4 Stochasticity in optimum phenotypic change

All of the simulations and calculations of *Δshift* in the above sections were deterministic. However, any recorded environmental parameter in nature, always fluctuates around a mean expectation over time (García Molinos and Donohue 2011). We thus wanted to assess whether adding stochasticity to the changes in the optimum affected our deterministic simulation results. We redid the numerical simulations of the above theoretical model but added stochasticity in the optimum phenotypic change (for stochasticity in the growth rate, see appendix A8). For details see appendix A3. We then assessed how *Δshift* performed under stochasticity in optimum phenotypic change and how the abovementioned factors affected *Δshift*

#### 2.2.5 Changing genetic variance

All of the above simulations were done assuming that genetic variation remained constant. This is true when a quantitative genetic trait was assumed to be controlled by infinite number of loci (Falconer and Mackay 1996). Here, we took into account the decreases in genetic variation that might occur due to directional change of the optimum phenotype. For an asexually reproducing population, the variance of the distribution of the breeding values was given by the equation (Bürger 2000):

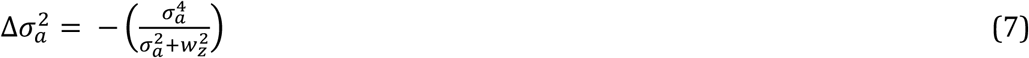

We used numerical iteration to solve the change in genetic variation over time using equation (7). We repeated all the simulations above and varied the levels of plasticity, reproductive rate, starting genetic variation, and environmental predictability, and calculated *Δshift* as before (appendix A4).

#### 2.5.6. EWS

We evaluated how shifts in EWS compared with shifts in mean trait in response to changes in the environment. First, we showed that our model exhibits non-catastrophic transcritical bifurcation (appendix A6). Next, we estimated standard deviation and autocorrelation at first-lag from abundance data as two main EWS indicators. Following this, shifts in EWSs and shifts in mean trait value were compared by calculating a metric called *Δshift_ews_* (appendix A6.4-A7). We evaluated *Δshift_ews_* for different levels of plasticity, genetic variation and reproductive rate. Furthermore, we modified our model to allow for catastrophic fold bifurcation by introducing an allee threshold at equation 2 (Dai et al. 2012; Hilker 2010). Similarly, for this fold-bifurcation model we evaluated *Δshift_ews_* (see appendix A6-A7 for details).

### 2.3 Experimental Data

In addition, we analyzed an experimental data where microcosm populations were forced to collapse by varying the rate of decline in food availability (Clements & Ozgul, 2016). Clements & Ozgul (2016) used replicate populations of protozoan ciliate *Didinium nasutum* that fed on *Paramecium caudatum*. In the experiment, four different treatments of rates of decline of prey availability were chosen: 1) fast; 2) medium; 3) slow, as well as a constant prey availability as the control treatment. A total of 60 replicate populations, 15 per treatment were used for the experiment. In our study, we used data only from the deteriorating environment treatments (i.e., fast, medium and slow decline in prey availability). We analyzed each population’s abundance and mean body size time series independently. We then calculated AUC and *Δshift* to qualitatively verify our theoretical simulation results (see appendix A5 for details).

## 3. Results

### 3.1 Analytical results

Before a shift in the optimum environment occurred, we assume that the population is perfectly adapted to its optimum phenotype 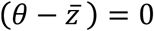. Considering the starting population size to be at K with no plasticity (b =0), equilibrium population size at any time *t* from equation (1), (3) and (4) is

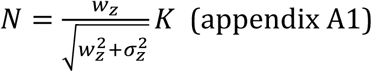

#### Case #1: Population declines before a trait shift

Let 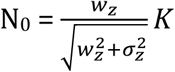 be the population size when the population is at its equilibrium. Next, when the environment changes by a large magnitude in one generation such that *δ* ≥ 1.5, the standardized population shift in one generation is (appendix A1):

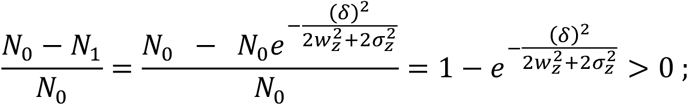

and, 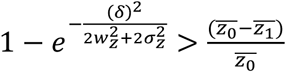

#### Case #2: Trait shifts before population decline

When the environment shifts by a small magnitude such that: *δ* ≤ 0.2 in one generation. The standardized population shift becomes (appendix A1):

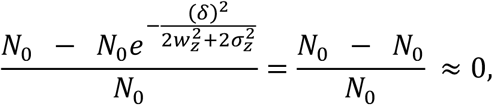

because, 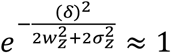 as *δ* is very low conditional to the fact that 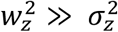. In this case, proportional shift in a trait is greater than zero and hence greater than the population shift (see appendix A1).

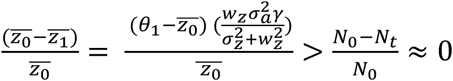

Our two extreme analytical cases showed that, if the optimum phenotype shifted by a large magnitude in just one generation, the decline in population size preceded shift in mean trait value immediately. However, this was not true in the scenario when the optimum shifts by a very small magnitude over the course of a single generation.

For the effect of adaptive plasticity see appendix A2.

### 3.2 Simulation results

#### 3.2.1 Environmental predictability

Higher predictability of the optimum phenotypic change caused *Δshift* to be positive when compared with lower predictability (Fig. 2). Low predictability of the optimum phenotypic change caused *Δshift* to be negative even when the optimum phenotype shifts by a magnitude of ~1.5 units in more than 15 generations (slow environmental shift) (Fig. 2).

**Figure 2.**
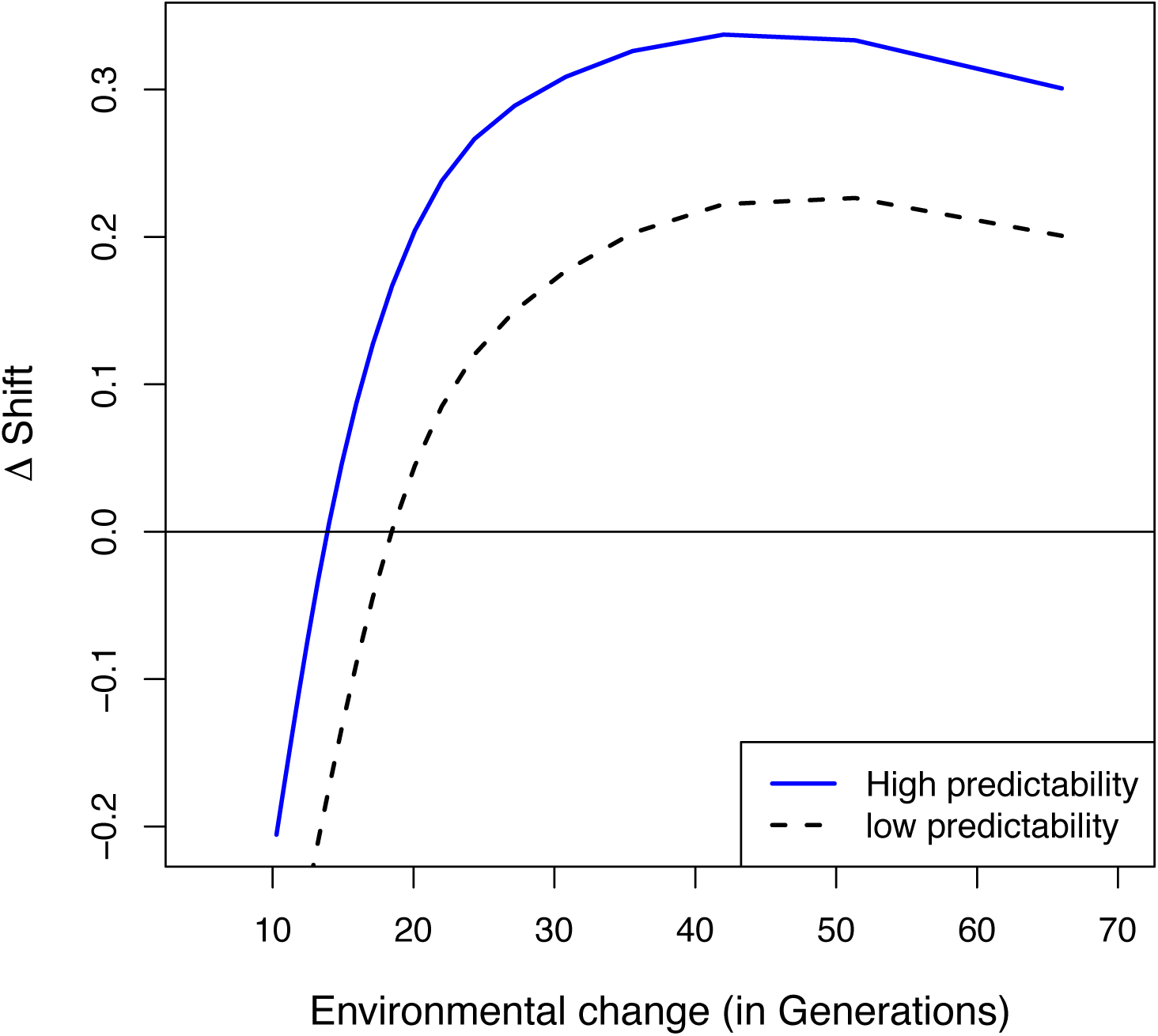
Effect of environmental predictability on the magnitude of *Δshift*, data from numerical simulations of the theoretical model. *Δshift* denotes how large and earlier the shift occurs in trait dynamics before a decline in the population. If *Δshift* is +ve, shift in average trait value precedes decline in population size and vice versa. The X-axis denotes the time in generations it takes for the optimum to change by a magnitude of 1.5 units, and hence 10 in the X-axis means the optimum shifts by a magnitude of 1.5 in 10 generations indicating a fast change and 60 means the optimum takes 60 generations to shift by a magnitude of 1.5 units indicating a very slow change. High environmental predictability (blue line] facilitates earlier occurrence of trait shift (*Δshift* being +ve] compared to when the environmental predictability is very low (dashed line]. *Δshift* is always −ve (which means population declines precede mean trait shifts] when the optimum environment shifts in less than or equal to 15 generations by 1.5 units. Parameters used: 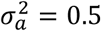; *R*_0_ = 1.2; *b* = 0.2; 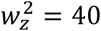.

#### 3.2.2 Genetic variation

Higher genetic variation 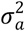 caused *Δshift* to be positive but this happened when the optimum environment shifted by a magnitude of 1.5 units in more than 15 generations (slow environmental shift) (Fig. 3C). If the optimum environment shifted to a new value by a magnitude of 1.5 units in less than 15 generations (fast shift), population size always declined before any visible shift in the trait value.

**Figure 3.**
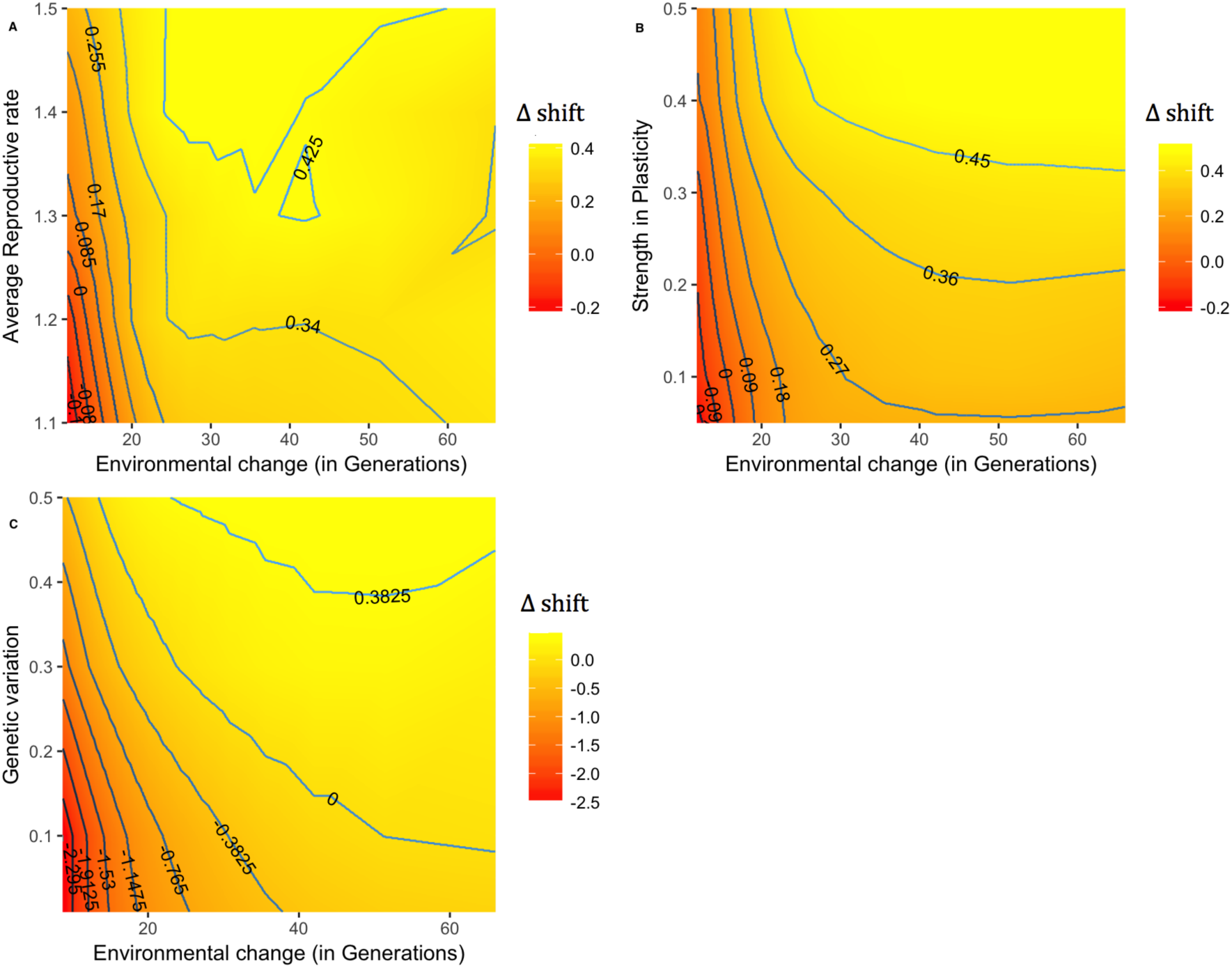
Contour plot showing the effect of net reproductive rate, plasticity and genetic variation on *Δshift* (contour lines]. Shown here are the results from numerical simulations of the theoretical model. For a trait shift to be informative of a population decline, the *Δshift* should be always greater than zero (contour line =0]. The X-axis denotes the time in generations it takes for the optimum to shift by a magnitude of 1.5 units.

#### 3.2.3 Adaptive plasticity

Values of plasticity (*b* > 0.2) caused *Δshift* to be positive (i.e., trait shifted earlier than population decline) even when the optimum shifted by a magnitude of 1.5 units in less than 15 generations (fast shift) (Fig. 3B). However, for lower values of plasticity (b <0.2), earlier shift in trait value would occur before the population declined only when the optimum environment shifted slowly (>15 generations) (Fig. 3B).

#### 3.2.4 Reproductive rate

For R_0_ of 1.2, trait shift occurred earlier than decline in population size i.e., *Δshift* > 0, given the optimum phenotype shifted slowly (by a magnitude of 1.5 units in ~15 generations or more). If the optimum phenotype shifted faster than that, average reproductive rate should have to be higher than 1.2 for *Δshift* > 0 and hence shift in the trait value would then be informative of a population decline (Fig. 3A).

#### 3.2.5 Stochastic change in the optimum phenotype

Higher 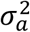, higher *b*, higher environmental predictability, and higher R_0_ substantially caused *Δshift* to be positive for this particular scenario (see appendix A3).

#### 3.2.6 Evolving genetic variation

In comparison to simulations with constant genetic variation, *Δshift* was smaller (less positive) for simulations where genetic variation evolved over time. In other words, *Δshift_constant variation_* > *Δshift_evolving variation_* (see appendix A4).

#### 3.2.7 EWS and shift in mean trait

We found that for both transcritical and fold-bifurcation models, shifts in mean trait could occur significantly before shifts in the two EWS indicators (Appendix A6-A7). This was particularly evident for medium to slow changes in the environment (Fig A13-A16).

### 3.3 Experimental results

Experimental data supported our analytical and simulation results qualitatively. Decline in prey availability from ~300 individuals of *P. caudatum* to zero happened over a period of 10, 15 and 20 days indicating three different rates: fast, medium and slow change in the environment respectively. Fast change in prey availability resulted in population decline preceding shift in mean body size in 7 out of 9 replicates (Fig. 4A). However, during medium rate of prey decline, shift in average body size preceded decline in population size in 7 out 9 replicate populations in and 4 out of 7 in slow decline in prey availability, qualitatively proving our analytical as well as our simulation results correct (Fig. 4A).

**Figure 4.**
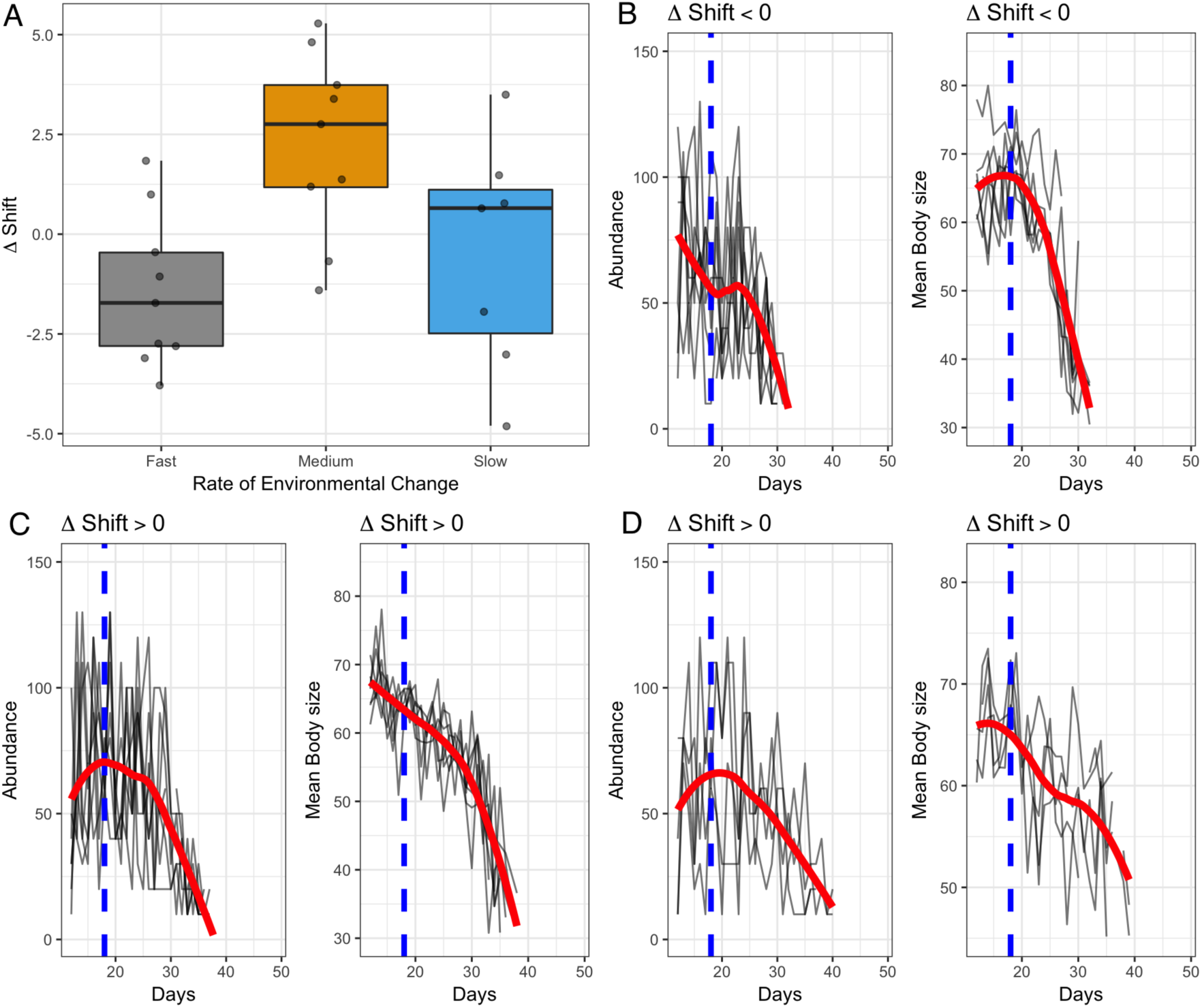
Ability of shift in body size dynamics to precede population decline (i.e., *Δshift*>0) under different environmental regimes, for the experimental data from Clements & Ozgul (2016). Note that *Δshift* < 0 if population decline precedes shift in mean body size and *Δshift* > 0 if shift in mean body size precedes population decline. (A) *Δshift* <0 for fast decline in prey availability indicating that population decline preceded shift in mean body size. For medium and slow decline in prey availability, *Δshift* was largely positive (*Δshift* >0). (B) Replicate abundance and body size time-series are shown for *Δshift* < 0 (fast decline in prey availability] and for *Δshift* >0 (C and D] (Medium decline and slow decline in prey availability respectively]. The blue dashed line in B, C and D indicates the starting point of environmental deterioration. The red line indicates *loess* smoothing across replicates.

## 4. Discussion

Predicting the fate of populations in response to environmental change is a key challenge in ecology. Recently developed methods have suggested that tracking fitness related traits might help achieve this goal (Clements and Ozgul 2016; Clements et al. 2017). Our analytical and simulation results suggested that when the optimum environment changes relatively slowly, shifts in average trait value may occur earlier than shifts in the population abundance. However, this was affected by the amount of genetic variation, strength in adaptive plasticity, environmental predictability, and speed of life history.

In our simulations, due to higher adaptive plasticity (b>0.3), shifts in the average trait value occurred before a decline in population size in response to a fast change in the optimum environment (i.e., shift of 1.5 magnitude by the optimum environment in less than 15 generations)(Fig. 3B, Fig. A2). Such a result was dependent on how predictable the environment was, as plastic response of the trait was mediated by how well the cue was related to a change in the optimum environment. If the cue was a function of the selecting environment, very high adaptive plasticity (b = 0.5) would then lead to a shift in average trait value before a decline in population size was observed even during a significant fast shift in the optimum phenotype (Fig. 3B, *b=0.5*).

While the positive influence of higher genetic variation on population persistence (Willi and Hoffmann 2009) and evolutionary rescue (Hufbauer et al. 2015; Gomulkiewicz, Krone, and Remien 2017) is relatively well studied, little is known about its transient effect on shifts in average trait value in response to changes in the optimum environment. Our result suggest that in response to a relatively slow directional change in the environment, high and constant genetic variation in a fitness-related trait will promote faster evolution in the average trait value and consequently will cause a faster shift in the average trait value before a decline in population size (Fig. 3C, Fig. A3). In addition, additive genetic variance is also expected to decrease over time due to directional selection acting on the trait (Barton and Keightley 2002). Here, in addition to the simulations with constant genetic variation, we carried out a set of simulations where we relaxed the assumption of constant genetic variation, and this slowed evolutionary change due to the depletion of genetic variation caused by directional selection (Fig. A6). For this reason, the magnitude of shift in average trait value was smaller and slower when compared with another trait shift under the assumption of constant genetic variation for the same scenarios of optimum environmental change (see appendix Fig. A6, A7). Besides directional selection, a decline in population size can also lead to a decrease in genetic variance (Ashander, Chevin, and Baskett 2016). Irrespective of the cause, low genetic variance will impede both evolutionary rescue (Hufbauer et al. 2015; Gomulkiewicz, Krone, and Remien 2017) as well as the predictability of population decline with the help of trait information.

Deterministic and stochastic modeling of population persistence in response to a changing environment had earlier revealed the positive affect of reproductive rate on influencing adaptation (Yvonne and Hoffmann 2009). Earlier studies have also reported that larger species tend to decline in population size more rapidly than smaller species (Olden, Hogan, and Zanden 2007; Collette et al. 2011) due to differences in life-history strategies and intrinsic growth rates between large and small organisms (Hutchings et al. 2012). Populations of fish with slow life histories (in the family Scombridae) declined faster than those which had comparatively faster speed of life history (Juan-Jordá et al. 2015). Our modeling results also reiterated that during a fast shift in the environment (shift of the optimum environment by 1.5 units in <15 generations), slow growing populations (R_0_ < 1.2) would decline before a shift in the average trait value could be observed (Fig. 3A). When we compared two populations with different reproductive rates, the magnitude of *Δshift* was found to be substantially larger for populations with higher R_0_ for a same rate of environmental shift (Fig. 3A, Fig. A9), due to the rapid declines of the populations with lower R_0_.

Our results also suggested that environmental predictability (the correlation between the cue and the optimum environment) was a key factor in determining the earlier occurrence of a shift in the average trait value in response to a change in the optimum environment. Such environmental predictability acted as an interactive factor, determining the speed and magnitude of shifts in trait value, which was driven particularly by the strength in plasticity. Earlier studies had also indicated the positive interactive effects of environmental predictability and adaptive plasticity on population dynamics (Reed et al. 2010; Ashander, Chevin, and Baskett 2016). Introducing stochasticity in the change in the optimum phenotype or in the growth rate (appendix A8) did not change our above results substantially (Fig. A4). In the case of stochastic change in the optimum phenotype, environmental predictability was particularly essential as the plastic response of the trait tracked the changes in the optimum which led to an earlier trait shift before the population declined (Fig.4B, Fig. A4).

EWS are shown to exist in models showing both non-catastrophic and catastrophic transitions (Kéfi et al. 2013). In relation to this, our results suggest that regardless of whether our model exhibited non-catastrophic transcritical or catastrophic fold bifurcation, shifts in mean trait value could occur before shifts in EWS. This was particularly evident for medium to slow change in the optimum environment (FigA10-A16). Such a shift in mean trait value occurring before EWS, was however slightly sensitive to variation in plasticity, genetic variation and net reproductive rate. Nevertheless, shift in mean trait value in conjunction with shift in EWS could be used as an indicator of imminent population declines (Clements and Ozgul 2016).

Our analytical and simulation results were qualitatively supported by experimental data (Fig. 4A). In both medium and slow decline in prey availability treatments, shifts in body size occurred before decline in population size. Shifts in body size in response to decline in prey availability that were seen in the experimental data could mostly be attributed to plasticity in body size, as the experimental population was clonal. During fast decline in prey availability, the plastic response of body size over time was not large enough to keep up with the pace of decline in prey availability and hence could not stabilize the loss in fitness. This led to rapid decline in population size before a significant body size shift. However, during medium decline in prey availability, plastic shift in body size was able to track the decline in prey availability in consequence of which a positive growth rate was maintained. However, since the decline in prey availability continued, the plastic capacity of body size in the population was depleted causing the population to eventually decline, which was later than shift in body size was observed. In case of slow decline in prey availability, the change was very slow that led to small plastic shifts in body size before a decline in abundance was seen. These small shifts in body size were not large enough in comparison to decline in abundance, which was reflected in some of the replicates. Shifts in body size thus could be an obvious indicator of environmental deterioration before a response in the population dynamics could be observed. Whether a trait could be considered as an additional indicator of how stressed a population is, would depend not only on the identity of the trait but also on the kind of environmental forcing.

The results that we presented here were specific to the parameter space, but were not restricted to any specific model system (see Chevin, Lande, and Mace 2010). In our model, we made two main assumptions. Firstly, adaptive plasticity in our trait remained constant and hence could not evolve. Studies have observed adaptive phenotypic change without being able to attribute it to genetic change (Ghalambor et al. 2007; Hendry, Farrugia, and Kinnison 2008). Our results, under the assumption of constant adaptive plasticity, still held since we explicitly dealt with transient dynamics after the optimum environment shifted. Secondly, we assumed linear reaction norms. Nonlinear reaction norms are modeled for secondary traits that are components of fitness, but such nonlinear reaction norms can evolve to linear ones in the long term (Gavrilets and Scheiner 1993).

In conclusion, we show that shifts in average trait value could precede shifts in EWS and population declines in response to a change in the optimum environment, and higher levels of genetic variation, adaptive plasticity, environmental predictability, and reproductive rate strengthened such an earlier shift in the average trait value. Using an experimental data we also showed that shifts in average body size could precede declines in population size, and hence could be indicative of a future population decline. Such a shift in mean body size preceding a decline in population size was possible if the change in the optimum environment was not fast relative to the generation time of the organism. Thus, shifts in traits may be useful for predicting population collapses in species where life histories are fast, the rate of change of the environment is relatively slow, and the environmental predictability is relatively high, giving hope that methods can be developed that for these signals in real world populations.

## 5. Acknowledgements

This research was supported by the European Research Council Grant #337785 and the Swiss National Science Foundation Grant #31003A_146445 to xx.

